# Rodent perirhinal cortex, but not ventral hippocampus, inhibition induces approach bias under object-based approach-avoidance conflict

**DOI:** 10.1101/2022.06.28.497792

**Authors:** Sandeep S. Dhawan, Carl Pinter, Andy C. H. Lee, Rutsuko Ito

**Affiliations:** Department of Psychology (Scarborough), University of Toronto, Toronto, ON M1C 1A4, Canada; Rotman Research Institute, Baycrest Centre, Toronto, ON M6A 2E1, Canada; Department of Cell and Systems Biology, University of Toronto, Toronto, ON M5S 3G5, Canada

**Keywords:** perirhinal cortex, decision-making, hippocampus, approach, avoidance, anxiety, behavioural inhibition, punishment, reward, optogenetics

## Abstract

Neural models of approach-avoidance (AA) conflict behavior and its dysfunction have focused traditionally on the hippocampus, with the assumption that this medial temporal lobe (MTL) structure plays a ubiquitous role in arbitrating AA conflict. We challenge this perspective by using three different AA behavioural tasks in conjunction with optogenetics, to demonstrate that a neighbouring region in rats, perirhinal cortex, is also critically involved but only when conflicting motivational values are associated with objects and not contextual information. The ventral hippocampus, in contrast, was found not to be essential for object-associated AA conflict, suggesting its preferential involvement in context-associated conflict. We propose that stimulus type can impact MTL involvement during AA conflict and that a more nuanced understanding of MTL contributions to impaired AA behaviour (e.g., anxiety) is required. These findings serve to expand upon the established functions of the perirhinal cortex while concurrently presenting innovative behavioural paradigms that permit the assessment of different facets of AA conflict behaviour.

## Introduction

Approach-avoidance (AA) conflict is elicited when an organism experiences competition between incompatible motivations of being attracted to and repelled by the same goal stimulus. Its effective resolution is vital for survival and everyday decision-making, while its dysfunction manifests across a spectrum of psychiatric disorders including anxiety and addiction (Aupperle and Paulus, 2010; Fricke and Vogel, 2020). Since the septo-hippocampal system was first theorised to mediate a behavioural inhibition system (BIS) that is activated to suppress approach responses under conflict situations (Gray and McNaughton, 2000), converging cross-species empirical evidence has highlighted a critical role for the ventral (rodent) or anterior (primate), but not dorsal/posterior portion of the hippocampus (HPC), in modulating anxiety and AA responding during high motivational conflict (Ito and Lee, 2016). For example, in addition to increasing anxiolytic behaviour in rodents, lesion or pharmacological inactivation of the ventral HPC (vHPC) increases approach behaviours to motivationally-conflicting learned stimuli (Bannerman et al., 2014, 2003, 2002; Schumacher et al., 2018, 2016; Yeates et al., 2019). Similarly, human functional magnetic resonance imaging (fMRI) and patient neuropsychological studies with analogous AA tasks have revealed anterior HPC involvement when participants experience high AA conflict (Bach et al., 2014; O’Neil et al., 2015).

Given the focus on the role of the HPC in AA conflict processing, very limited empirical work has explored the potential contributions of the surrounding medial temporal lobe (MTL) cortices. This is somewhat surprising given the intricate anatomical and functional relationships between the MTL structures. Moreover, a revised formulation of the BIS postulated the involvement of the entorhinal cortex and perirhinal cortex (PRC) in the detection and resolution of AA conflict (Gray and McNaughton, 2000), a suggestion that has not yet, to our knowledge, been fully examined empirically. Theoretical models of MTL function posit that stimulus type can impact MTL structure recruitment during cognition. Specifically, the HPC is implicated in scene and context-based processing, while the PRC is predominantly involved in object-associated processes, both within the domain of memory and even beyond (Graham et al., 2010; Murray et al., 2007; Zeidman and Maguire, 2016). Since ethological anxiety tasks and paradigms of AA conflict processing have typically employed spatial stimuli (e.g., scene images) or required environmental exploration (e.g., maze navigation) (Bach et al., 2014; Bannerman et al., 2002; O’Neil et al., 2015; Schumacher et al., 2018, 2016), it raises the question of whether the HPC plays a ubiquitous role in AA conflict processing across all stimulus domains or whether other MTL structures may also play a critical role depending on the stimuli involved.

Here we designed two rodent AA tasks using objects as target stimuli to test the hypothesis that differential MTL recruitment occurs during AA conflict processing in a stimulus-type specific manner in rodents. In both tasks, rats were trained to associate object-pairings with either appetitive or aversive outcomes and were subsequently presented with these objects in a conflict arrangement while the PRC or vHPC (CA3 subfield) was optogenetically inhibited. However, each task was conducted in a different apparatus, specifically a runway or shuttle box (Figure 1), to elicit a different type of avoidance behaviour (passive vs. active). The PRC was also inhibited while animals completed an established ‘contextual’ AA task, known to be dependent on the vHPC (Schumacher et al., 2018, 2016; Yeates et al., 2019). Across both object-based tasks, we observed that inhibition of rodent PRC, but not the vHPC, induced a significant approach bias in response to object-associated motivational conflict. In contrast, there was no impact of PRC inhibition when conflict was represented by contextual cues. These findings implicate a hitherto unconsidered substrate in the arbitration of AA conflict; they emphasize the need to consider the involvement of the broader MTL in AA conflict processing and have implications for our understanding of the neural substrates underlying disorders in which AA conflict behaviour is compromised.

**Figure 1.**
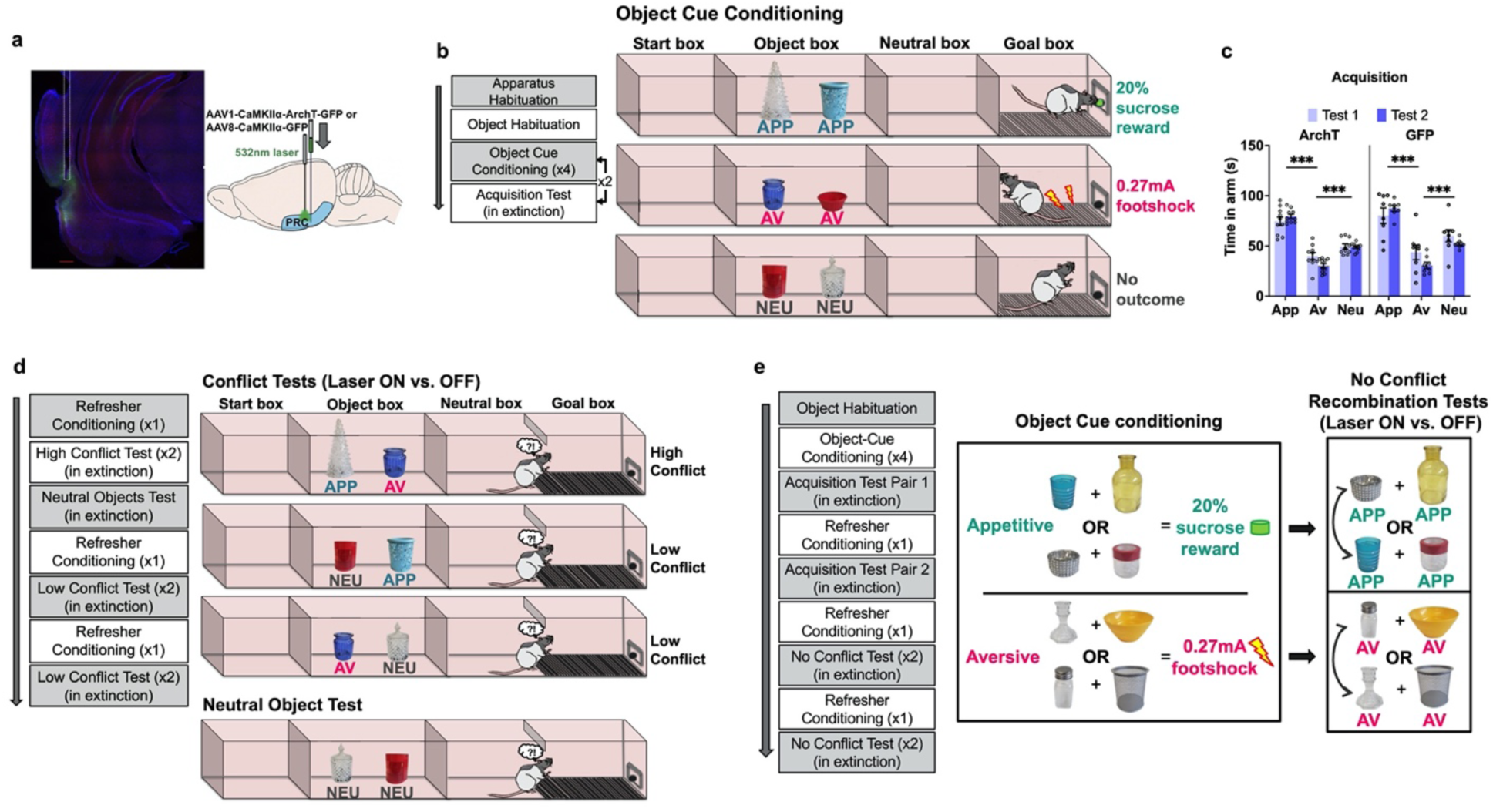
Object AA conflict runway task. **(a-b)** After injection with ArchT or GFP in the PRC, rats were put through object cue conditioning to learn the outcomes associated with appetitive (APP), aversive (AV) and neutral (NEU) object pairs. **(c)** Acquisition data indicated successful learning in both groups (*** p < 0.001). **(d)** Rats then underwent a series of tests in extinction, specifically high conflict (APP + AV), neutral (NEU + NEU), and low conflict (APP + NEU or AV + NEU). **(e)** A no conflict recombination task was then administered, in which rats first learned a new set of object pairs and were then presented with recombined object pairs composed of objects of the same valence.

## Results

### Optogenetic inhibition of PRC increases approach behaviour to motivational conflict represented by objects in a runway paradigm

Rats were first surgically infused with inhibitory archaerhodopsin T (ArchT) or green fluorescent protein (GFP - control) and implanted bilaterally with optic fibres into the PRC. Rats were then trained on a novel behavioural paradigm in which they first acquired the incentive values of 3 pairs of objects (appetitive [sucrose], aversive [electric shock], and neutral [no outcome] pairs) in a customised runway apparatus comprising 4 successive compartments: a start box; an object box containing two objects; a neutral box in which the rat was held temporarily after the object box; and finally, a goal box, in which the associated outcomes were delivered during training (Figure 1c). Objects were selected to be visually and texturally distinct from one another since PRC is implicated in the discrimination of objects with overlapping features (Murray et al., 2007). Acquisition of object valences was assessed after four (test 1) and eight (test 2) conditioning sessions, without laser treatment. PRC rats acquired the object-outcome associations successfully by test 2, with rats spending the most time in the goal box after exposure to the appetitive object pair (p < 0.001), and the least time after aversive object pair exposure (p < 0.001), compared to goal box time in neutral trials (valence: F(2, 32) = 329.64, p < 0.001; valence x test: F(2, 32) = 11.97, p < 0.001) (Figure 1d). The pattern of valence acquisition was comparable between ArchT and GFP animals (construct: F(1, 16) = 1.84, p = 0.19; valence x construct: F(2, 32) = 1.29, p = 0.29; test x construct: F(2, 32) = 0.22, p = 0.64).

Animals were then administered a series of tests in which recombinations of the learned objects were presented to elicit a high (appetitive-aversive) or low (appetitive-neutral; aversive-neutral) level of motivational conflict, or the original neutral object pair was presented (Figure 1d). When animals were exposed to a high conflict pairing, optogenetic inhibition of PRC (laser on) significantly increased time spent in the goal box compared with trials completed without inhibition (laser off), and compared to GFP control animals with laser on and off (laser x construct: F(1, 16) = 29.03, p < 0.001; all post-hoc: p < 0.001) (Figure 2a), indicative of an increase in approach behaviour under motivational conflict. Furthermore, PRC inhibition led to a significant decrease in the number of retreats from the goal box (laser x construct: F(1,16) = 16.80, p < 0.001), although there was no effect on the number of goal box entries (laser x construct: F(1,16) = 1.92, p = 0.19) (Figure 2b) during the choice period. PRC inhibition also had no impact on the latency to enter each of the object, neutral and goal boxes (laser x construct: F(1,16) = 2.45, p = 0.14; laser x construct x box = F(1, 32) = 1.41, p = 0.26) (Supplementary Figure 1a). Collectively, these results indicate that when faced with high motivational conflict, PRC-inhibited rats entered the goal box as readily as control animals but stayed longer and retreated less, indicative of a potentiated approach bias under conflict. This effect was not observed in the neutral object test in which PRC inhibition had no effect on the time spent in the goal box (laser x construct: F(1,14) = 1.32, p = 0.27) (Figure 2c).

**Figure 2.**
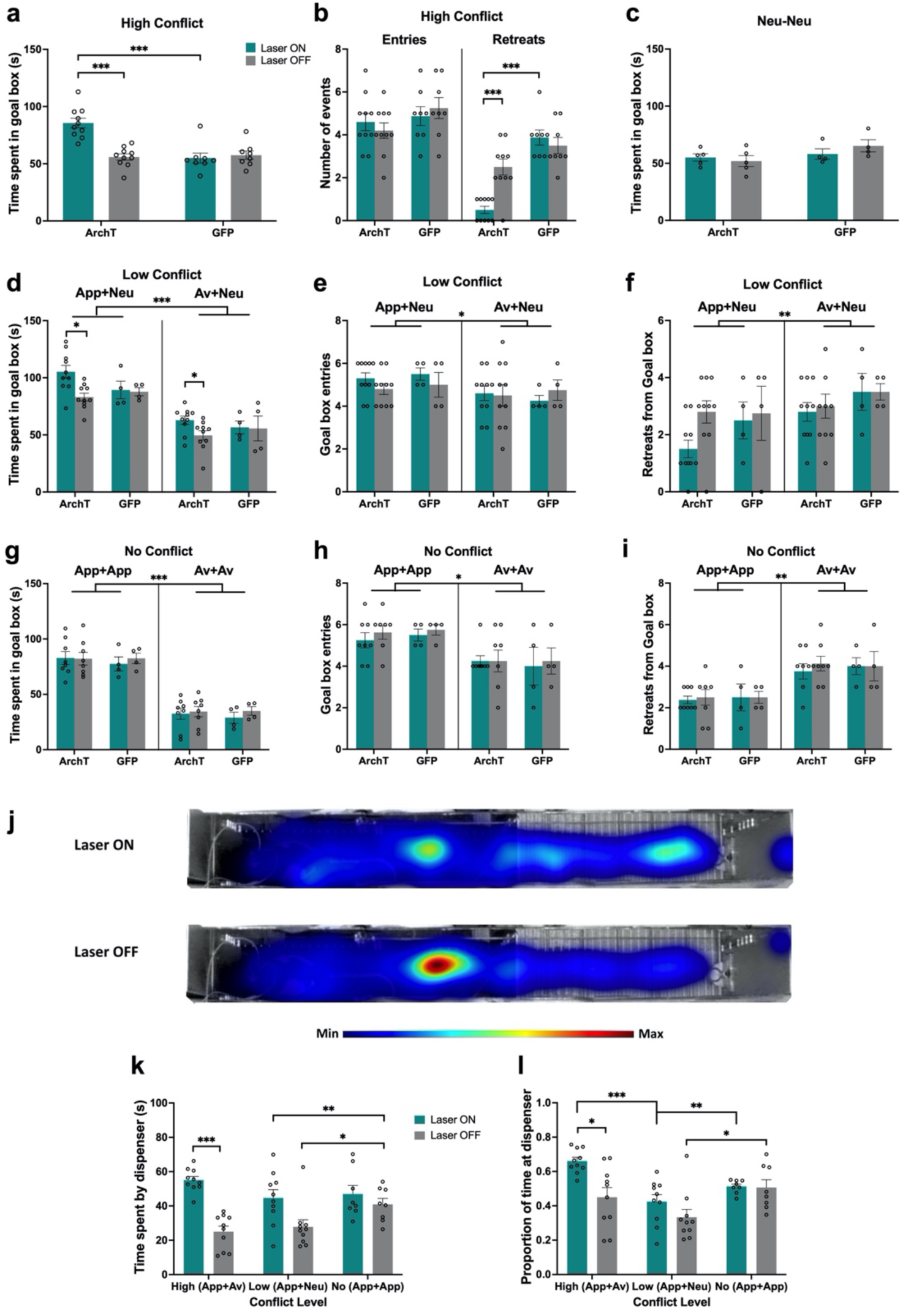
Impact of PRC inhibition on object AA conflict runway task performance. **(a-b)** PRC inhibition significantly increased time spent in the goal box and reduced the number of retreats in the high conflict test. **(c)** There was no effect of PRC inhibition on AA behaviour in the neutral test. **(d)** Similar to the high conflict test, PRC inhibition increased time spent in the goal box in both low conflict tests. **(e-f)** PRC inhibition did not impact the number of entries or retreats in the App + Neu or Av + Neu low conflict tests, although there was a main effect of valence, with a greater number of entries for App + Neu and more retreats for Av + Neu. **(g-i)** PRC inhibition did not affect AA behaviour on the no conflict tests. A main effect of valence was observed for all measures, indicating intact valence retrieval for the recombined test pairs. **(j)** Heatmap plots for the high conflict test. **(h-i)** PRC inhibited rats spent more time by the sucrose dispenser, as measured by total time or proportion of time, in the high and low conflict tests but not the no conflict test. *** p < 0.001, ** p < 0.01, * p < 0.05

Following a refresher conditioning session, we next investigated whether PRC-inhibited animals showed altered conflict processing when faced with a lower degree of motivational conflict (Figure 1e). PRC inhibition increased the amount of time animals spent in the goal box after exposure to both appetitive-neutral and aversive-neutral object pairs (laser x construct: F(1,12) = 4.88, p = 0.047) (Figure 2d) but had no effect on the number of entries into (laser x construct: F(1,12) = 0.60, p = 0.45), nor retreats from (laser x construct: F(1,12) = 0.24, p = 0.64) the goal box during the choice period (Figure2e-f). PRC inhibition also had no impact on the latency to enter each of the runway boxes (laser x construct: F(1,12) = 0.19, p = 0.67; laser x construct x box: F(2, 24) = 0.91, p = 0.42) regardless of the valence of the object pairing (valence: F(1,12) = 2.38, p = 0.149) (Supplementary Figure 1b-c).

Notably, PRC inhibition did not impair the ability to discriminate appetitive and aversive valences (valence: F(1,12) = 109.10, p < 0.001; laser x valence x construct: F(1,12) = 0.62, p = 0.45), with all animals spending significantly more time in the goal box for appetitive-neutral pairings than aversive-neutral (p < 0.001, Figure 2d). Furthermore, all PRC rats made fewer entries (p = 0.050) and exhibited more retreat behaviour (p = 0.009) when presented with an aversive-neutral pairing compared with an appetitive-neutral pairing (Figure 2e-f). This indicates that valence recall was intact in PRC inhibited rats and is also consistent with the findings from the neutral object pair test (Figure 2c). Thus, it is the presence of motivational conflict, whether low or high, which led to increased approach behaviour following PRC inhibition.

To rule out the possibility that the observed effects of PRC inhibition were driven by a failure to respond to novel recombinations of object pairings rather than conflict processing, we repeated the runway task in which animals were trained to associate a new set of 4 object pairs with either appetitive or aversive outcomes (2 pairs each) (Figure 1e). Animals acquired the cue-outcome associations successfully by the first test, and spent significantly more time in the goal box after exposure to the appetitive object pairs compared with the aversive pairs, with comparable acquisition between sets of objects pairs within each valence (valence: F(1, 13) = 240.69, p < 0.001; pairing: F(1, 10) = 0.12, p = 0.74; valence x pairing: F(1, 10) = 0.007, p = 0.94) (Supplementary Figure 1d). The pattern of object-cue acquisition was comparable between ArchT and GFP animals (construct: F(1, 10) = 0.96, p = 0.35; pairing x construct: F(1, 10) = 0.58, p = 0.46). Rats were then administered a within-valence ‘no conflict’ recombination test (appetitive-appetitive; aversive-aversive). PRC inhibition did not lead to significant changes in the time spent in the goal box after exposure to novel appetitive or aversive object pairs (laser: F(1, 10) = 0.60, p = 0.46; laser x construct: F(1, 10) = 0.37, p = 0.55) (Figure 2g). Furthermore, both ArchT and GFP animals could readily discriminate between valences during this test (valence: F(1,10) = 186.80, p < 0.001; valence x construct: F(1, 10) = 0.02, p = 0.88). PRC inhibition also had no effect on the number of entries into (laser x construct: F(1,10) = 0.007, p = 0.93) nor retreats from (laser x construct: F(1,10) = 0.15, p = 0.71) the goal box, and furthermore, all PRC-inhibited rats made fewer entries (p = 0.01) and exhibited more retreat behaviour (p = 0.003) when presented with an aversive pairing compared with an appetitive pairing (Figure 2h-i).

Finally, to investigate whether animals exhibiting approach bias spent an increased amount of time in the vicinity of the sucrose dispenser in expectation of reward, we analysed the amount of time and proportion of total goal box time that ArchT animals spent in a demarcated ‘dispenser zone’ (final 18 cm or 1/3 of goal box) during the high, low and no conflict test sessions. When faced with high conflict, PRC inhibition led to animals spending significantly more time by the sucrose dispenser compared to high conflict test sessions without optogenetic inhibition (p < 0.0001), whereas an increase in time spent by the dispenser elicited by low conflict stimuli approached significance (p = 0.052; laser: F(1, 18) = 24.34, p < 0.0001; laser x conflict: F(2, 32) = 5.43, p = 0.0093) (Figure 2j-k). As expected, in the absence of PRC inhibition, animals spent more time by the dispenser during no conflict trials compared to high conflict (p = 0.002) and low conflict (p = 0.03) trials.

When considering the time spent by the dispenser as a proportion of the total time in the goal box, PRC-inhibited rats spent a higher proportion of time during high conflict trials compared with low conflict (p = 0.0008) and no conflict (p = 0.0012) trials, as well as high conflict trials completed with the laser off (p = 0.015; laser: F(1, 18) = 7.8, p = 0.012, conflict: F(1.79, 28.65) = 11.19, p = 0.00038 laser x conflict: F(2, 32) = 3.34, p = 0.048) (Figure 2l). Laser off animals also spent proportionally more time by the dispenser during no conflict trials compared with low conflict trials (p = 0.017), with no difference between no conflict and high conflict trials (p = 0.89).

Collectively, these findings indicate that the increase in approach behaviour during the high and low conflict tests arises from PRC-mediated impairments in conflict processing, rather than impairments in valence retrieval or novel stimulus recombination processing. Furthermore, in the absence of the normal PRC functioning, animals spend more time by the sucrose dispenser perhaps reflecting a greater anticipation for a reward outcome.

### Optogenetic inhibition of PRC decreases avoidance behaviour to motivational conflict represented by objects in a shuttle box paradigm

To investigate whether the observed increase in goal box preference time exhibited by PRC-inhibited rats was specific to the conditions elicited by the runway task, the same PRC rats completed another novel behavioural paradigm conducted in a modified two-way active avoidance ‘shuttle box’ apparatus (Figure 3a-c). In the first phase of the paradigm, appetitive or aversive object-cue pairs used in the ‘no conflict’ recombination test in the runway task were placed in opposite ends of a two-compartment apparatus, behind transparent barriers. In each trial, animals were allowed to visually sample the objects for 1 min, after which the transparent barrier was raised to allow animals access to the associated outcome (sucrose or shock). At the same time, a central guillotine door dividing the two compartments was raised to give rats the opportunity to shuttle into the opposite compartment to escape the shock outcome associated with the aversive object cue pair. Acquisition of escape behaviour was assessed after four (test 1) and eight (test 2) conditioning sessions, without laser treatment. PRC rats demonstrated the expected behaviour by test 2, with rats exhibiting significantly shorter escape latencies after exposure to the aversive object pair compared with exposure to the appetitive object pair (valence: F(1, 9) = 22.23, p = 0.0011; valence x test: F(1, 9) = 7.81, p = 0.021) (Figure 3d). The pattern of escape acquisition was comparable between ArchT and GFP animals (construct: F(1, 9) = 3.07, p = 0.11; valence x construct: F(1, 9) = 3.01, p = 0.12; test x construct: F(1, 9) = 0.86, p = 0.38). PRC rats also spent more in the outcome-associated area (stay behavior) when exposed to the appetitive object pair compared with the aversive object pair (valence: F(1, 9) = 232.94, p < 0.0001; valence x test: F(1, 9) = 9.84, p = 0.012) (Figure 3e). This pattern of stay behavior was also comparable between ArchT and GFP animals (construct: F(1, 9) = 2.66, p = 0.13; valence x construct: F(1, 9) = 4.31, p = 0.07; test x construct: F(1, 9) = 0.81, p = 0.39).

**Figure 3.**
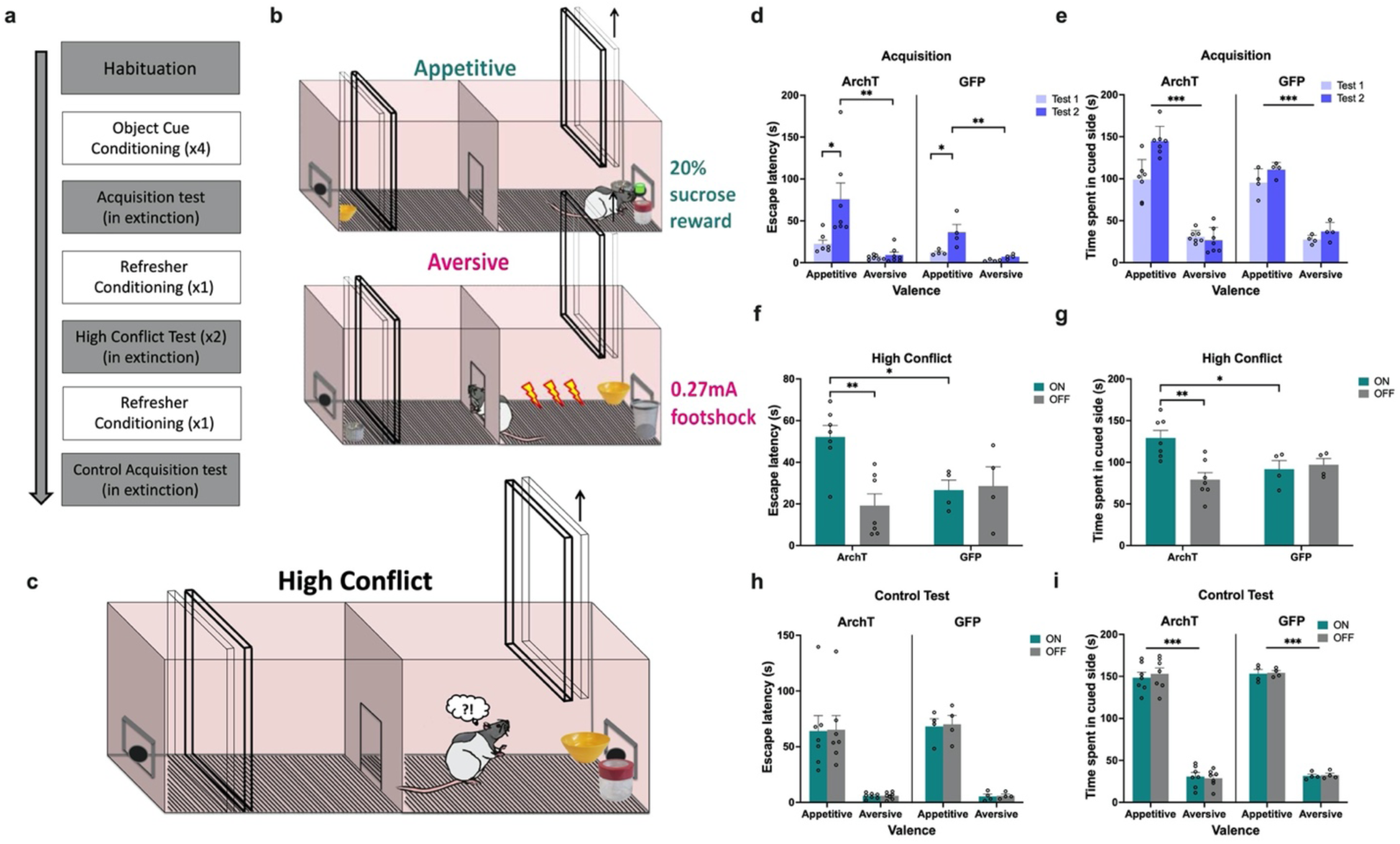
Impact of PRC inhibition on object AA conflict shuttle box task performance. **(a)** Timeline of paradigm. **(b)** Rats first learned to stay to receive a reward when exposed to an appetitive object pair, and to escape to avoid footshock when exposed to an aversive object pair. **(c)** Rats were then exposed to a high conflict object pairing in extinction. **(d-e)** All rats demonstrated intact acquisition of AA behaviour. **(f-g)** PRC inhibition led to an increased escape latency in the high conflict test and a greater amount of time spent in the cued side of the shuttle box. **(h-i)** PRC inhibition did not impact escape latency or time spent in the cued side in a subsequent control no conflict test. *** p < 0.001, ** p < 0.01, * p < 0.05

When animals were next exposed to a high conflict pairing, optogenetic inhibition of PRC (laser on) significantly prolonged escape latencies compared with trials completed without inhibition (laser off; p < 0.005), and compared to GFP control animals (p < 0.05) with laser on and off (laser x construct: F(1, 9) = 10.39, p = 0.010) (Figure 3f), indicative of a decrease in avoidance behaviour under motivational conflict. Furthermore, despite making a comparable number of re-entries into the outcome-associate area during the test session (laser x construct: F(1, 9) = 1.57, p = 0.24), PRC-inhibited animals spent significantly longer in the outcome-associated area containing the conflict object pair compared with trials completed without inhibition (laser off; p<0.005), and compared to GFP control animals (p < 0.05) with laser on and off (laser x construct: F(1, 9) = 6.80, p = 0.028)(Figure 3g).

Following ‘refresher conditioning,’ PRC-rats completed a set of control tests, in which conditioned cue acquisition tests with and without PRC inhibition were completed. PRC-inhibition had no effect on the duration of escape latencies for either appetitive or aversive valences (laser: F(1, 9) = 0.34, p = 0.58; laser x construct: F(1, 9) = 0.03, p = 0.86; valence x construct: F(1, 9) = 0.06, p = 0.81), with all rats exhibiting significantly shorter escape latencies for aversive than appetitive trials (valence: F(1, 9) = 41.88, p < 0.001; construct: F(1, 9) = 0.06, p = 0.82) (Figure 3h). Similarly, PRC-inhibition had no effect on the amount of time spent in the cued side of the apparatus (laser: F(1, 9) = 0.26, p = 0.62; laser x construct: F(1, 9) = 0.01, p = 0.94; valence x construct: F(1, 9) = <0.01, p = 0.96), with all rats spending significantly less time in the cued side during aversive than appetitive trials (valence: F(1, 9) = 322.94, p < 0.0001; construct: F(1, 9) = 0.63, p = 0.45) (Figure 3i). Collectively, these results suggest that PRC-inhibition led to both a decrease in avoidance behaviour and an increase in approach/stay behaviour under motivational conflict, and that PRC-inhibited animals could readily discriminate between appetitive and aversive object cues.

### PRC inhibition does not impact contextual AA conflict processing

To investigate whether the observed alteration in object-based AA conflict processing extended to conflict represented by ‘context-like’ cues, we administered a radial arm maze (RAM) AA task that is vHPC-dependent, with large vHPC lesions and ventral CA3 (vCA3) or dentate gyrus inactivation increasing approach behaviour in the face of high AA conflict (Schumacher et al., 2018, 2016; Yeates et al., 2019).

Rats first learned the valences of three visuotactile cues (appetitive, aversive, and neutral) that spanned the length of three different maze arms (Figure 4a). Analysis of time spent in a given arm after four (test 1) and eight (test 2) conditioning sessions, without laser treatment, revealed that all rats acquired the cue-outcome associations successfully by test 2 (valence: F(2, 32) = 146.02, p < 0.0001; valence x test: F(2, 32) = 24.26, p < 0.0001), regardless of construct (construct: F(1,16) = 0.17, p = 0.69; test x construct: F(1,16) = 0.01, p = 0.92; valence x construct: F(2, 32) = 1.51, p = 0.24) by spending significantly more time in the appetitive arm (both tests p < 0.001) and less time in the aversive arm (both p ≤ 0.003) compared with the neutral arm (Figure 4b).

**Figure 4.**
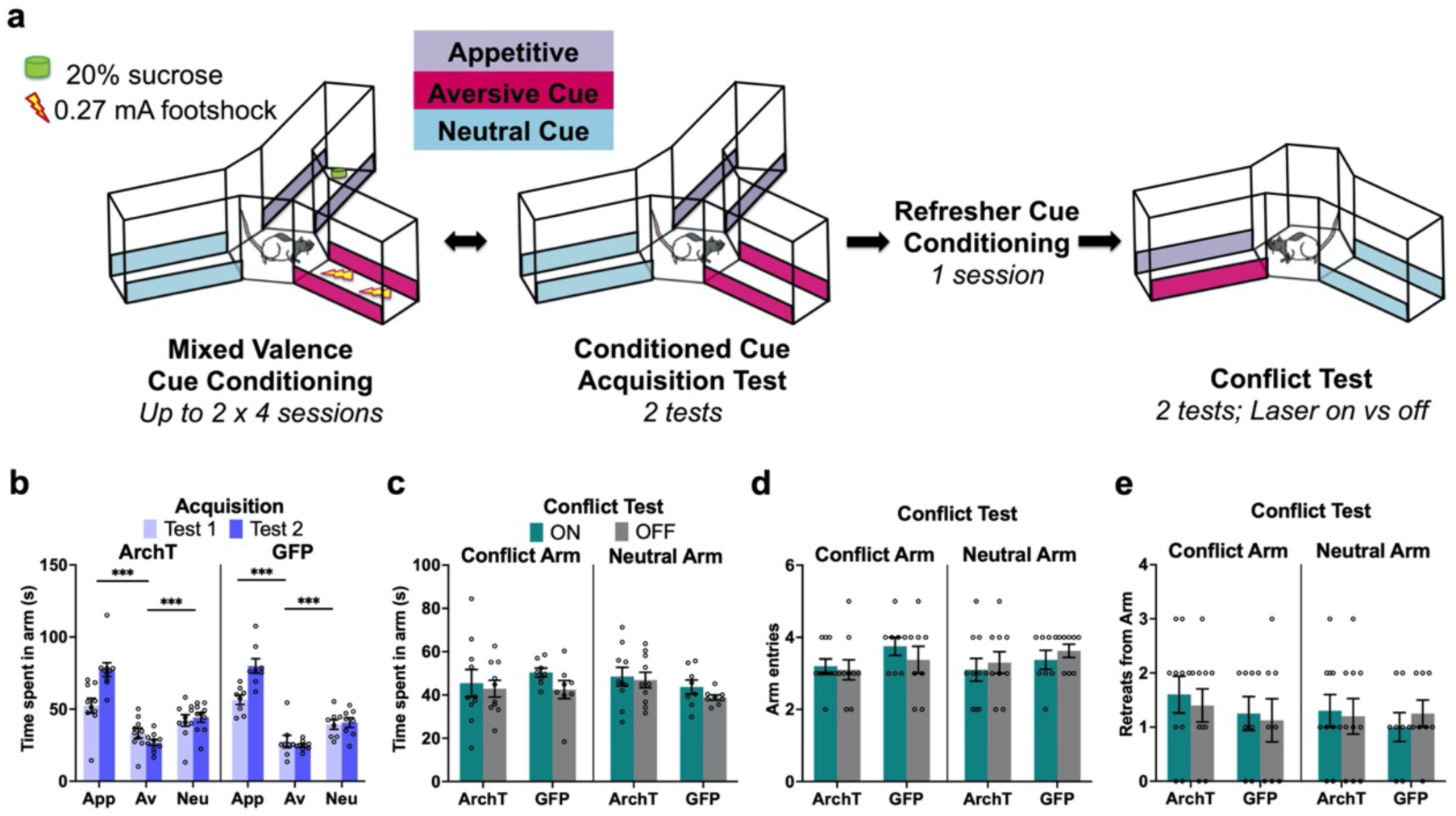
No effect of PRC inhibition on contextual AA conflict behaviour. **(a)** PRC rats underwent a contextual AA task known to be vHPC-dependent, in which they first learned the outcomes associated with appetitive, aversive, and neural cues and then underwent a conflict test in extinction. **(b)** Both ArchT and GFP rats demonstrated successful valence acquisition. **(c-e)** PRC inhibition had no effect on choice behaviour during the conflict test.

Rats then completed two AA conflict tests during which they could freely explore between a conflict arm containing a combination of appetitive and aversive cues, and a neutral arm containing the neutral cue. PRC inhibition did not significantly change the time spent in either arm (laser: F(1,16) = 4.06, p = 0.06; laser x construct: F(1,16) = 1.06, p = 0.32) (Figure 4c), and both ArchT and GFP animals spent a comparable duration of time in both the conflict and neutral arms (arm: F(1,16) = 0.09, p = 0.77; construct: F(1,16) = 0.31, p = 0.58; arm x construct: F(1,16) = 2.47, p = 0.14). Moreover, PRC inhibition did not significantly alter the number of entries made into or retreats from either the conflict or neutral arms (entries: laser: F(1,16) = 0.001, p = 0.98; construct: F(1,16) = 3.36, p = 0.09; laser x construct: F(1,16) = 0.07, p = 0.8; *retreats*: laser: F(1,16) = 0.03, p = 0.86; construct: F(1,16) = 0.5, p = 0.49; laser x construct: F(1,16) = 0.2, p = 0.66) (Figure 4d-e). The number of entries and retreats also did not differ between conflict and neutral arms (*entries:* arm: F(1,16) = 0.001, p = 0.98; *retreats:* arm: F(1,16) = 1.19, p = 0.29,) for both ArchT and GFP animals (*entries*: arm x construct F(1,16) = 0.07, p = 0.8; *retreats*: arm x construct: F(1,16) = 0.43, p = 0.52). Thus, while PRC is critical for AA conflict processing associated with discrete objects, it may play a minimal role in contextual AA conflict.

### Optogenetic inhibition of vCA3 does not affect object-associated AA conflict processing

To investigate whether the vHPC plays a role in object-associated AA conflict processing, rats with either ArchT or GFP and optical fibre implants in the vCA3 (Figure 5a) were administered the object runway paradigm. vCA3 rats demonstrated successful valence learning across the two acquisition tests (valence: F(2, 26) = 160.44, p < 0.0001; valence x test: F(2,26) = 23.73, p < 0.0001) (Figure 5b), and at test 2 spent the most time in the goal box after appetitive object exposure and the least time after aversive object exposure in comparison to neutral trials (both p < 0.001). Object valence acquisition was similar for ArchT and GFP animals (construct: F(1, 13) = 3.85, p = 0.07; valence x construct: F(2, 26) = 0.20, p = 0.82) with the exception that ArchT rats spent less time in the goal box compared to GFP rats during acquisition test 1 (test x construct: F(1,13) = 7.13, p = 0.019; test 1 construct: F(1,13) = 10.94, p = 0.006). ArchT and GFP animals did not, however, differ at the end of training (test 2 construct: F(1,13) = 2.21, p = 0.16).

**Figure 5.**
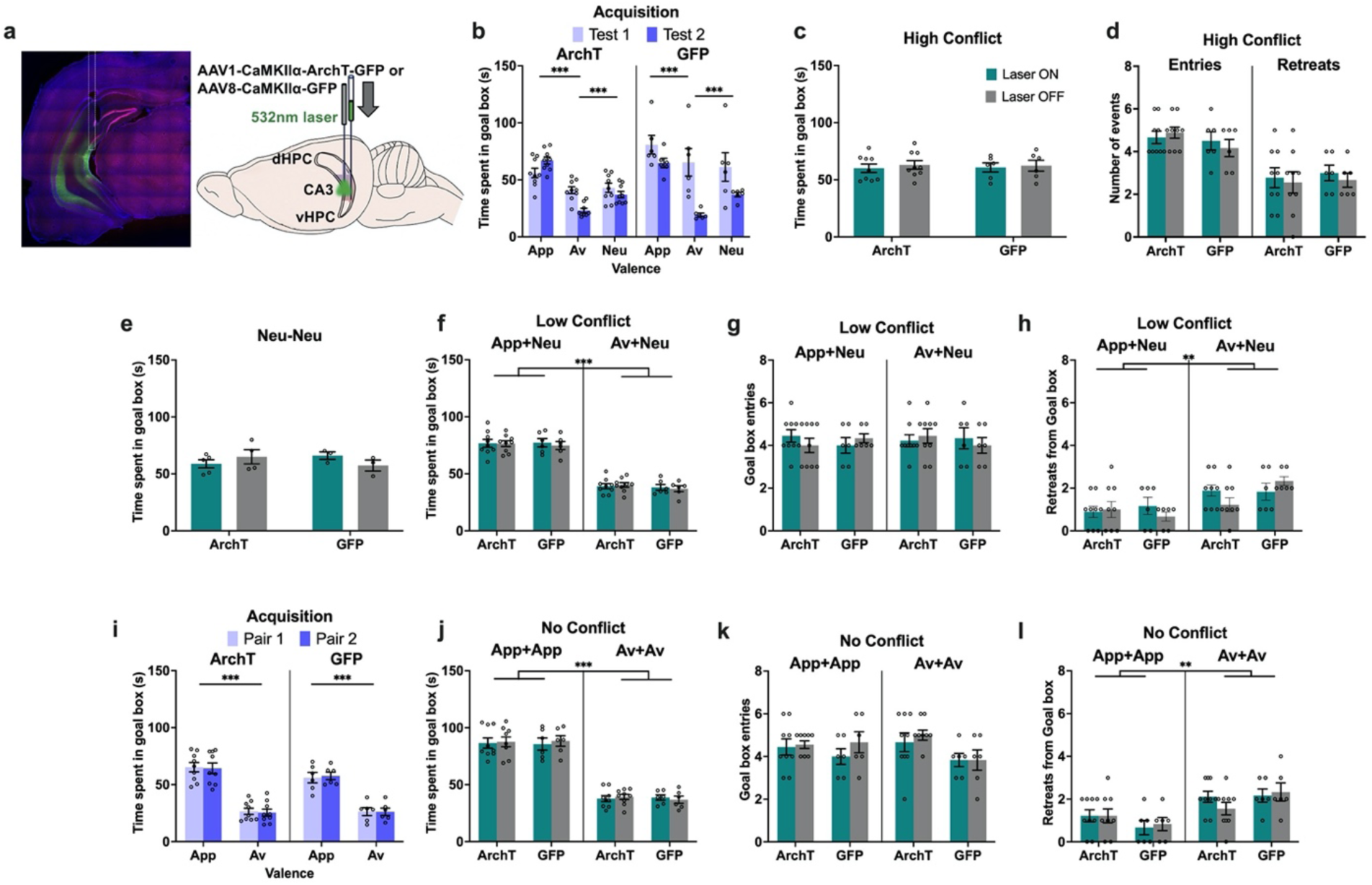
No effect of vCA3 inhibition on object AA conflict runway task performance. **(a)** Rats injected with ArchT or GFP in the vCA3 underwent the object runway task. **(b)** Both groups learned the appetitive (App), aversive (Av) and neutral (Neu) object pairs successfully. **(c-h)** vCA3 inhibition did not impact any behavioural measure on the high conflict, neutral, or low conflict recombination tests. **(i)** Rats successfully learned a new set of object pairs for the no conflict recombination tests. **(j-l)** vCA3 inhibition did not impact performance on the no conflict recombination tests. *** p < 0.001, ** p < 0.01

In contrast to PRC inhibition, there was no effect of vCA3 inhibition on time spent in the goal box in the high conflict test session (laser: F(1, 13) = 2.92, p = 0.11; construct: F(1,13) = 0.001, p = 0.99; laser x construct: F(1, 13) = 0.93, p = 0.60) (Figure 5c). Indeed, an overall comparison across regions revealed that the impact of laser manipulation was significantly different between PRC and vCA3 animals (laser x region x construct: F(1, 29) = 22.83, p < 0.0001). vCA3 inhibition also did not impact other behavioral measures (Figure 5d) including the number of retreats (laser x construct: F(1, 13) = 0.03, p = 0.86), entries made into the goal box (laser x construct: F(1,13) = 0.58, p = 0.46) and latency to enter (laser x construct: F(1,13) = 0.37, p = 0.55). vCA3 inhibition also did not affect time spent in the goal box when rodents were presented with the neutral object pair (laser: F(1,11) = 0.03, p = 0.90; laser x construct: F(1,11) = 2.36, p = 0.15) (Figure 5e).

Similar to the high conflict test, vCA3 inhibition did not impact the time spent in the goal box during the low conflict test (laser: F(1,13) = 0.11, p = 0.75; laser x construct: F(1,13) = 0.29, p = 0.60) (Figure 5f) and crucially, there was a significant difference between vCA3 and PRC rats when compared directly (laser x area x construct: F(1, 25) = 5.13, p = 0.032). All vCA3 animals could readily discriminate between valences as reflected by time spent in the goal box (valence: F(1,13) = 550.18, p < 0.0001; valence x construct: F(1,13) = 0.25, p = 0.63) and retreats (valence: F(1,13) = 13.80, p = 0.003). There was, however, no effect of vCA3 inhibition on entries into and retreats from the goal box (both valence x construct: F(1,13) ≤ 0.55, p ≥ 0.47) (Figure 5g-h).

Following successful acquisition of a new set of appetitive and aversive object pairs (valence: F(1,13) = 361.37, p < 0.0001; pairing: F(1, 13) = 0.006, p = 0.94; valence x pairing: F(1, 13) = 0.19, p = 0.67; pairing x construct: F(1, 13) = 0.14, p = 0.72) (Figure 5i), vCA3 inhibition also did not impact time spent in the goal box in the within-valence no conflict recombination test (laser: F(1,13) = 0.14, p = 0.72, ηp^2^ = 0.01; laser x construct: F(1,13) = 0.04, p = 0.85) (Figure 5j), with entries into and retreats from the goal box also unaffected (both valence x construct: F(1,13) ≤ 1.10, p ≥ 0.31) (Figure 5k-l). Both ArchT and GFP animals could discriminate between the two valences by spending more time in the goal box and retreating less for appetitive pairs compared to aversive pairs (valence: F(1,13) = 481.57, p < 0.0001; valence x construct: F(1,13) = 0.03, p = 0.88). In sum, vCA3 inhibition had no impact on any aspect of performance on the object AA runway task.

vCA3 rats also completed the object-based shuttle box, and all rats demonstrated successful establishment of an active avoidance response toward aversive, but not appetitive object pairings, across two acquisition tests, exhibiting significantly shorter escape latencies after exposure to the aversive object pair compared with exposure to the appetitive object pair (valence: F(1, 13) = 119.71, p < 0.0001; valence x test: F(1,13) = 92.12, p < 0.0001; construct: F(1, 13) = 1.43, p = 0.25; valence x construct: F(1, 13) = 1.18, p = 0.3; test x construct: F(1, 13) = 0.03, p = 0.87) (Figure 6a). All vCA3 rats also spent more in the outcome-associated area when exposed to the appetitive object pair compared with the aversive object pair (valence: F(1, 13) = 384.84, p < 0.0001; valence x test: F(1, 13) = 9.41, p = 0.0090; construct: F(1, 13) = 0.01, p = 0.98; valence x construct: F(1, 13) = 0.68, p = 0.42; test x construct: F(1, 13) = 1.28, p = 0.28) (Figure 6b).

**Figure 6.**
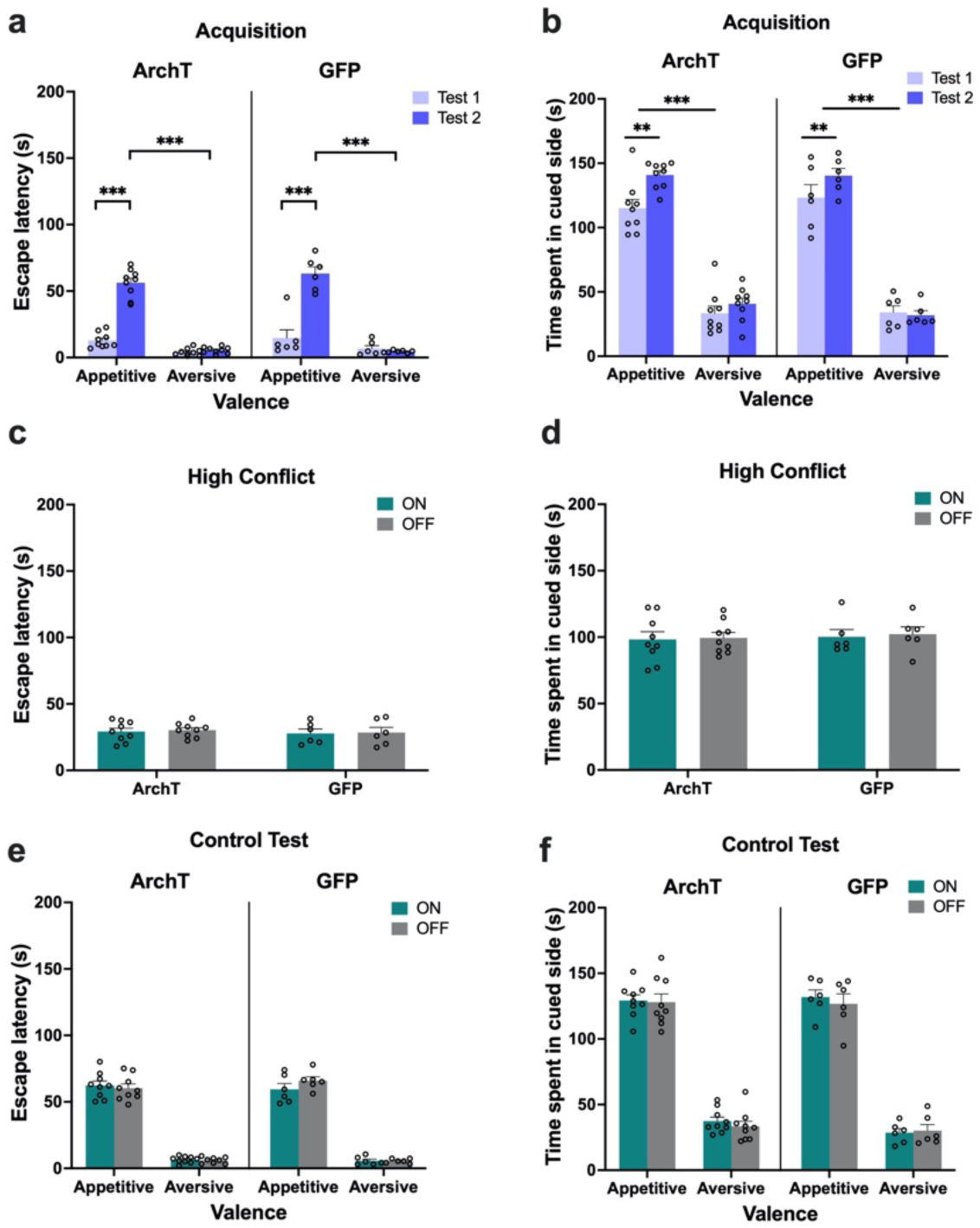
No impact of vHPC inhibition on object AA conflict shuttle box task performance. **(a-b)** Both ArchT and GFP vCA3 rats demonstrated intact acquisition of AA behaviour on the shuttle box task. **(c-d)** vCA3 inhibition had no effect on escape latency or time spent in the cued side in the high conflict test. **(e-f)** vCA3 inhibition also did not impact behaviour on the control no conflict test. *** p < 0.001, ** p < 0.01

In contrast to PRC rats, there was no effect of vCA3 inhibition on escape latency behaviour when exposed to the high conflict object pairing (laser: F(1,13) = 0.48, p = 0.5; construct: F(1,13) 0.03, p = 0.87; laser x construct: F(1,13) = 0.29, p = 0.87) (Figure 6c). An overall comparison across regions revealed that the impact of laser manipulation was significantly different between PRC and vCA3 animals (laser x region x construct: F(1, 22) = 13.78, p = 0.0012). vCA3 inhibition also had no effect on the time spent in the outcome-associated area containing the conflict object pair (laser: F(1,13) = 0.26, p = 0.62; construct: F(1,13) 0.12, p = 0.74; laser x construct: F(1,13) = 0.02, p = 0.89) (Figure 6d). Similarly, there was a significant difference between vCA3 and PRC animals when compared directly (laser x region x construct: F(1, 22) = 7.98, p = 0.010).

vCA3 inhibition also had no effect on the duration of escape latencies for either appetitive or aversive valences during the control acquisition tests (laser: F(1, 13) = 0.51, p = 0.49; laser x construct: F(1, 13) = 3.13, p = 0.1; valence x construct: F(1, 13) = 0.25, p = 0.63), with all rats exhibiting significantly shorter escape latencies for aversive than appetitive trials (valence: F(1, 13) = 826.68, p < 0.0001; construct: F(1, 13) = 0.03, p = 0.88) (Figure 6e). Similarly, vCA3-inhibition had no effect on the amount of time spent in the cued side of the apparatus (laser: F(1, 13) = 0.86, p = 0.37; laser x construct: F(1, 13) = 0.04, p = 0.84; valence x construct: F(1, 13) = 0.69, p = 0.42), with all rats spending significantly less time in the cued side during aversive than appetitive trials (valence: F(1, 13) = 565.77, p < 0.0001; construct: F(1, 13) = 0.3, p = 0.59) (Figure 6f).

In sum, these results indicate that vCA3-inhibition had no impact on object-associated motivational behaviour whether under approach-avoidance conflict or the presence of appetitive or aversive cues alone.

### Optogenetic inhibition of the PRC and vCA3 reduces cFos+ expression

To confirm optogenetic inhibition of cellular activity in the PRC and vCA3, cFos immunohistochemistry was used with a PRC-dependent task: novel object recognition (NOR), in which rats explored an open maze containing a novel and a familiar object; or a vHPC-dependent task: the elevated plus maze (EPM), in which rats explored two anxiogenic open arms and two ‘safe’ closed arms.

For the NOR task, a discrimination ratio (difference between novel and familiar object exploration divided by total exploration) revealed that laser-on PRC-ArchT animals exhibited a significant novel object recognition impairment (laser: F(1, 14) = 12.53, p = 0.003, construct: F(1, 14) = 6.86, p = 0.02, laser x construct: F(1, 14) = 10.84, p < 0.0001) (Figure 7a) compared with laser-off Arch T and PRC-GFP control animals (all p ≤ 0.001). In the EPM, laser-treated vCA3-ArchT animals spent a significantly increased proportion of time in the open arms (laser: F(1, 11) = 8.02, p = 0.003 construct: F(1, 11) = 14.42, p = 0.003; laser x construct: F(1, 11) = 7.1, p = 0.022 compared with laser-off Arch T and vCA3-GFP control animals (all p ≤ 0.001) (Figure 7b).

**Figure 7.**
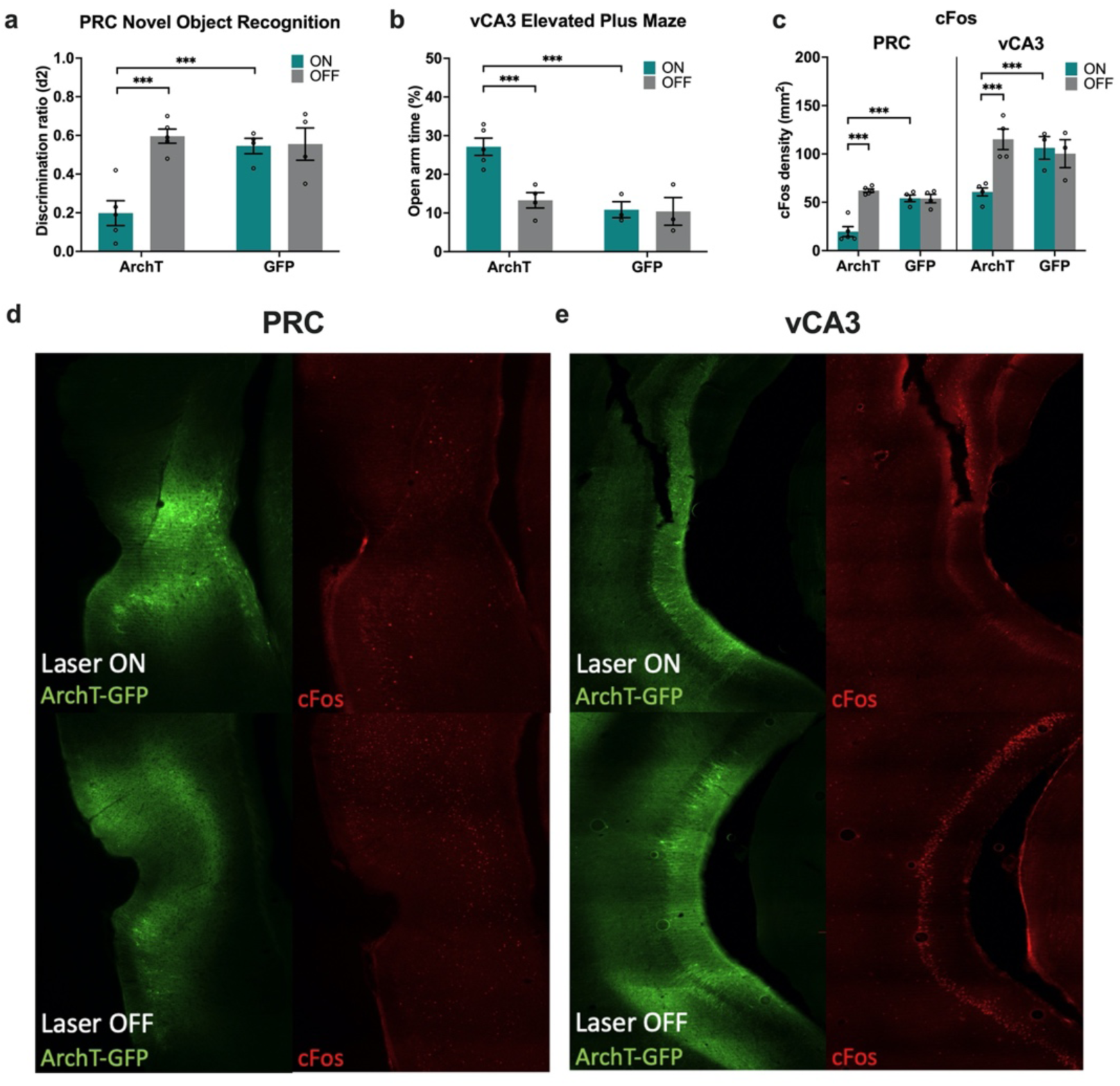
Effect of optogenetic inhibition on PRC- or vCA3-dependent control tasks and cFos expression. **(a)** PRC inhibition disrupted the ability of rats to discriminate a novel and familiar object on the novel object recognition task. **(b)** vCA3 inhibition increased time spent in the open arm of the elevated plus maze. **(c-e)** PRC and vCA3 inhibition was associated with decreased cFos levels in each respective area. *** p < 0.001

Histology revealed that all animals presented with robust GFP expression within the PRC or vCA3, with optic fiber implantation immediately dorsal to the viral site. ArchT animals that completed either the NOR (PRC) or EPM (vCA3) tasks with the laser on prior to sacrifice demonstrated a significant reduction in cFos labelling in the PRC (laser: F(1, 14) = 29, p < 0.0001; construct: F(1, 14) = 11.14, p = 0.005; laser x construct: F(1, 14) = 29.69, p < 0.0001) or vCA3 (laser: F(1, 11)= 5.81, p = 0.035; construct: F(1, 11) = 3.77, p = 0.07; laser x construct: F(1, 11) = 12.78, p = 0.004), compared with ArchT animals that completed the tasks with laser off and GFP-control animals (all p ≤ 0.001) (Figure 7c-e). Thus, laser treatment in ArchT animals significantly reduced neural activation.

## Discussion

Using a set of original object-based AA paradigms, we found that optogenetic inhibition of the rodent PRC, but not the vCA3, resulted in both a robust increase in approach behaviour and decrease in avoidance behaviour during motivational conflict elicited by the presentation of discrete object pairs associated with unmatching affective values. In contrast, PRC inhibition did not disrupt behaviour during a contextual vHPC-dependent AA task. Critically, PRC-inhibition did not disrupt AA behaviour in response to neutral stimuli or novel re-configurations of objects with the same valence, suggesting that the observed impact of PRC inhibition on object-associated AA conflict processing was unlikely to be driven by impairments in mnemonic functioning.

Our finding that optogenetic inhibition of PRC, but not vCA3, disrupted object-associated motivational conflict behaviour contrasts with a plethora of studies detailing a role for the rodent vHPC in the resolution of AA conflict (Ito and Lee, 2016). However, a common characteristic of these studies is the involvement of spatial/contextual stimuli, for example, AA conflict associated with spatial locations in ethological tests of anxiety or contextual cues with learned incentive values, with broad vHPC damage or subfield-specific inactivation of the vCA3 and DG leading to increased approach behaviour, and vCA1 inactivation leading to greater avoidance (Bannerman et al., 2003, 2002; Schumacher et al., 2018, 2016; Yeates et al., 2019). The present study advances this work by utilising discrete objects as target stimuli and raises the possibility that the vHPC may not play a ubiquitous role in AA conflict processing. Indeed, an absence of anterior HPC involvement has been previously reported in an instrumental approach-avoidance decision task in non-human primates (Wallis et al., 2019). Additionally, we recently reported significant involvement of human PRC rather than HPC activity, as measured by fMRI, during the resolution of AA conflict induced by objects with opposing valences (Chu et al., 2021). The current findings not only provide invaluable reverse-translational evidence for the rodent PRC mediating analogous behaviour, but uniquely demonstrate the necessity of PRC to the arbitration of object-associated AA conflict. Importantly, our observation that PRC inhibition did not alter behaviour towards a contextual representation of motivational conflict (i.e. conflicting ‘context-like’ bar cues that span a radial maze arm) underlines the specificity of the reported PRC effects to object stimuli, and aligns with theoretical viewpoints that posit a degree of stimulus specificity across MTL structures (Davachi et al., 2006; Graham et al., 2010; Murray et al., 2007; Zeidman and Maguire, 2016).

Although the observed involvement of the PRC in the arbitration of AA conflict may, at first sight, be unexpected given that this area is traditionally associated with mnemonic function, its contribution to motivational processes is plausible considering its reciprocal connectivity with limbic centres of the brain, including the amygdala and orbitofrontal, prelimbic and infralimbic cortices (Agster et al., 2016; Agster and Burwell, 2009; Burwell and Amaral, 1998; Pereira et al., 2016). The non-human primate PRC has been implicated in reward-related behaviour in a role that goes beyond the basic processing of stimulus-valence associations and may be crucial for mediating more nuanced relationships between cues, behaviour, and reward outcome (Suzuki and Naya, 2014). Specifically, while PRC damage/inactivation does not eliminate reward-related behaviour, it does impact an animal’s ability to track and flexibly adapt to changes in the magnitude and schedules of reinforcement of expected reward associated with visual stimuli (Clark et al., 2013; Liu et al., 2000; Liu and Richmond, 2000). Moreover, electrophysiological data point towards a role for PRC in processing configurations of multiple visual stimuli to predict upcoming reward (Ohyama et al., 2012). Our data complement and extend these findings by suggesting that rodent PRC is not simply necessary for the retrieval of object valence, as evidenced by valence-appropriate discriminative responding for low conflict (i.e. approaching more for appetitive-neutral vs. aversive-neutral), neutral and same valence object pairs during PRC inhibition, and may be engaged as part of the BIS to suppress approach when a violation of expected reward/punishment contingencies (e.g., conflict) is detected. Indeed, the absence of a functional PRC led to animals spending a greater proportion of time by the sucrose dispenser when presented with high conflict stimuli in the runway task, compared with stimuli with low or no motivational conflict, reflecting greater approach in anticipation of impending reward. Similarly, PRC-inhibited animals exhibited increased approach, and decreased avoidance of motivationally conflicting objects in the shuttle box task. Our finding of PRC, but not vHPC involvement in modulating conflict-elicited AA behaviour challenges the perspective proposed by Gray and McNaughton (2000) that MTL areas such as the PRC and entorhinal cortex are recruited to provide stimulus information of conflicting goals to the HPC, and that the HPC engages in resolving response conflicts. The present data would suggest that the PRC is an integral component of the BIS when conflicting configurations of learned object stimuli are encountered, independently of HPC recruitment.

An intriguing question is why PRC inhibition potentiated approach, as opposed to avoidance, of an uncertain outcome. In the present study, the excitatory/principal neuronal population (i.e., CaMKIIα-expressing cells) in PRC was targeted and silenced. These neurons receive powerful excitatory inputs from the basolateral amygdala, lateral amygdala, and medial prefrontal cortex (Curtis and Pare, 2004; Sah et al., 2003; Smith and Pare, 1994), which can override a strongly inhibitory intrinsic network within PRC (Kajiwara and Tominaga, 2021), and increase the likelihood of signal propagation to the entorhinal cortex and HPC (Kajiwara et al., 2003; Paz and Pare, 2013). In contrast, stimulation of local, short range neocortical inputs to PRC have been shown to evoke inhibitory potentials, suggestive of the recruitment of inhibitory interneurons giving rise to feedforward inhibition (Martina et al., 2001; Unal et al., 2013). Thus, the absence of glutamatergic projection neurons in the PRC could conceivably disrupt the balance of excitation/inhibition in PRC, and lead to a loss of output to a downstream target that is critical in suppressing approach in the face of motivational conflict. In the absence of an effect of inhibiting vHPC, a prominent projection target of PRC, on the expression of object-based AA conflict, we propose the nucleus accumbens, or septum as alternative candidate downstream areas that subserve this function. These areas receive direct projections from PRC (Agster et al., 2016) and are intrinsically organised (due to GABAergic principal neurons) to inhibit areas further downstream that are concerned with the execution of motor programs, and motivated behaviour (Floresco, 2015; Wirtshafter and Wilson, 2021).

Finally, given the well-established role of the PRC in recognition memory and the processing of novelty signals (Brown and Aggleton, 2001; Winters et al., 2008), it is important to highlight that the present effects of PRC inhibition cannot be explained by a disruption to this function. PRC intact rats typically increase exploration of novel compared to familiar stimuli (Winters et al., 2008) and yet here, PRC inhibition did not change choice behaviour during a ‘no-conflict’ test in which novel, recombined stimulus pairs within the same valence were presented. Similarly, it is also unlikely that the observed PRC-mediated impairments stem from a deficit in the perceptual processing of the two objects that comprised the high and low conflict pairs, as evidenced by the ability of PRC-inhibited animals to discriminate between valences during the low conflict test. Furthermore, although there is a delay between stimulus presentation and outcome presentation in the object-based runway AA task, it is unlikely that impairments in trace conditioning, in which the PRC have been implicated (Kholodar-Smith et al., 2008), can account for the PRC-mediated impairments in conflict processing. When stimulus presentation occurred in the outcome-associated area in the object-based shuttle box task (i.e., no delay between stimulus and outcome presentation), a similar increase in approach behaviour towards high conflict object pairs was observed.

To conclude, we demonstrate that rodent PRC is involved in the resolution of AA conflict, particularly when the goal stimulus is object-based. Thus, stimulus type may fundamentally alter how the brain represents motivational conflict, which may in turn recruit differential MTL structures for the purpose of AA conflict resolution. Our findings also have implications for our understanding of mental disorders in which AA behaviour is awry and suggest that a singular focus within the MTL on HPC dysfunction may be inadequate. Further work is needed on delineating the network of structures involved in various aspects of the conflict process, along with examining how a changing stimulus gradient (e.g., object vs. context vs. social) may recruit different brain structures to resolve conflict.

## Methods

### Subjects

Subjects were 33 male Long Evans rats (Charles River Laboratories, QC, Canada) weighing between 320 and 380g at the start of the experiment. Following surgery, rats were individually housed to prevent damage to the implanted optic fiber, and maintained at a constant room temperature of 22°C under a 12h light/dark cycle (lights on at 7:00am). Water was available *ad libitum*, and animals were fed *ad libitum* up until food-motivated experimental training commenced. Food-restricted animals were maintained at 85-90% of their baseline free-feeding weight. All experiments were conducted during the light cycle and were done in accordance with the regulations of the Canadian Council of Animal Care and approved by the University and Local Animal Care Committee of the University of Toronto.

### Surgery

Rats received viral infusions to either the PRC (*n* = 18; *n* = 10 ArchT, *n* = 8 GFP controls) or the vCA3 region of the HPC (*n* = 15; *n* = 9 ArchT, *n* = 6 GFP controls). Rats were anesthetized with isoflurane (Benson Medical, ON, Canada) and secured in a stereotaxic frame (Steolting Co, IL) with the incisor bar set at –3.3mm below the interaural line. An incision along the midline of the skull was made and the fascia retracted by small skin clips to reveal bregma. For PRC surgeries, the temporalis muscle was carefully peeled back from the temporal ridge to access lateral injection sites. Small burr holes were created directly over the injection sites using a dental drill, and 0.5µl of virus (either AAV1-CaMKIIa-ArchT-GFP or AAV8-CaMKIIa-GFP, Addgene, MA) was infused bilaterally with a 1 µl Hamilton syringe into either the PRC (AP: –5.2mm, ML: ±6.7mm, DV: – 7.3mm) or the vCA3 (AP: –5.2mm, ML: ±4.8mm, DV: –7.0mm) over 5 min, with the needle left *in situ* for a further 5 min. Optic fibers were then implanted 0.5mm ventral to the infusion site, and were secured in place with dental cement anchored to jeweler screws implanted in the skull. Rats were given at least 7 d to recover in their home cages before beginning behavioural experiments.

### Apparatuses

#### Object-based Approach-Avoidance Task apparatus

Testing was conducted in three radial arm maze arms [each 77 cm (L) X 11.5 cm (W) X 35 cm (H)] arranged to form a continuous ‘runway’ [231 cm (L)], with the entire apparatus wrapped in red cellophane to minimize reliance on extra-maze cues. The runway was segmented into 4 compartments (start box, object box, neutral box, and goal box) which were separable by stainless-steel guillotine door inserts. The entire stainless-steel grid floor was covered in opaque black Plexiglas, except for the goal box, which contained exposed grid flooring connected to a shock generator (Med Associates, VT, USA). A stainless-steel dish connected to a sucrose dispenser was presented at the end of the grid flooring (in the goal box), which the animals were required to traverse in order to obtain reward.

#### Object-based Shuttle box apparatus

Behavioural testing was conducted in a modified active avoidance ‘shuttle box’ apparatus [54 cm (L) X 26 cm (W) X 32.5 cm (H)] (Coulbourn Instruments, Whitehall, PA), which was divided into 2 equally-halved compartments [27 cm (L) X 26 cm (W) X 32.5 cm (H)] separated by a central guillotine door, with the entire apparatus wrapped in red cellophane to minimize reliance on extra-maze cues. Two removable panels [40 cm (L) X 25 cm (W)], one opaque and one transparent, were inserted in each side of the apparatus, which served to prevent recognition of and access to the object pairs, respectively. The transparent panel contained 2 rows of holes (each 4 cm apart), positioned at the rat’s eye level, to permit olfactory sampling. Two stainless-steel dishes connected to sucrose dispensers were presented at opposing ends of the apparatus, located behind both panels.

#### Radial Arm Maze (RAM) Approach-Avoidance Task apparatus

Behavioural testing was conducted in a three-arm radial maze apparatus, as previously described (Nguyen et al., 2019; Schumacher et al., 2018, 2016; Yeates et al., 2019). Briefly, each arm [50 cm (L) X 11.5 cm (W) X 35 cm (H)] was connected to a hexagonal central hub compartment [11.5 cm (W) X 35 cm (H)], with each arm wrapped in red cellophane to minimize reliance on extra-maze cues and arranged 120° relative to the adjacent arm. Arm entrances were blocked by stainless-steel guillotine doors. The grid flooring for each arm was connected to a shock generator (Med Associates, VT) and led to a stainless-steel dish connected to a sucrose dispenser.

#### Novel Object Recognition (NOR) apparatus

Behavioural testing was conducted in a transparent open-field apparatus [50 cm (L) X 50 cm (W) X 50 cm (H)], wrapped with black plastic to reduce anxiety. The apparatus was lined with home-cage bedding, which was replaced on days of testing. The bedding was agitated after each trial to eliminate potential odour traces.

### Stimuli

#### Object cues

Given the established role for the PRC in the resolution of feature ambiguity between highly similar objects (Saksida and Bussey, 2010), we attempted to maximize the similarity between selected objects by restricting the height (from 3.5–20 cm) and composition (either glass, plastic or metal) of the object-pair. A collection of ‘junk objects’ (i.e., no prior reinforcement history and no natural significance to the rats) were obtained for behavioural testing for the three behavioural experiments (Object-based Runway task, Object-based Shuttle box task, NOR). For NOR task, the pairs of objects were composed of the same material so they could not be readily discriminated by olfactory cues, whereas object exploration preference was used to assign pairings for the object-based AA tasks (see task procedure 2.5.3 below). The objects were attached to the apparatus floor with a hook-and-loop fastener. All objects were cleaned with 70% ethanol solution after each trial for all experiments.

#### Bar cues

A set of visuotactile cues were used in the RAM task, which consisted of wood panel inserts [46.5 cm (L) X 9.6 cm (H)] affixed to the length of each arm with hook-and-loop fasteners. Three sets of bar cues were used, with two sets wrapped in either duct tape or a denim cloth material, and the last set being an unwrapped, varnished wooden material. Bar cues were wiped down with a 70% ethanol solution between trials.

### Behavioural Procedures

PRC-rats and VCA3-rats first completed testing in the object-based AA runway task (Figure 1). They then underwent testing in the object-based AA shuttle box task. The PRC-rats were then trained in the RAM AA task, prior to completing a final NOR test before euthanasia. vCA3-rats were administered a final elevated plus maze task before euthanasia.

#### Object-based Approach-Avoidance Runway Task

##### Habituation

All rats were given 4 days of habituation to permit exposure to the runway apparatus and the object cues. On day 1, rats were confined to the start box for 30s, and then permitted to explore the entire apparatus for 3.5 min. On Day 2, following a 30s confinement to the start box, rats could proceed into and were confined within the object box (without any objects) for 1 min, with re-entry to the start box prohibited by a stainless-steel guillotine door. Rats could then proceed to the neutral box, in which they were confined for 1 min, with re-entry to the object boxes prohibited. Following this, they were confined to the goal box for 30s before being removed from the apparatus. Days 3 & 4 introduced the rats to the 6 object-cues used for the object-based AA task; each rat performed 3 daily trials, with each trial presenting 2 object-cues, and thus rats were exposed to all 6 object-cues daily. The order that the objects appeared for each rat was counterbalanced between animals and habituation sessions, and the time spent exploring the objects was recorded. Object habituation began with 30s confinement to the start box, and rats were then permitted 1 min to explore the 2 object-cues during a given session. Rats then entered the neutral box, in which they were confined for 1 min, followed by the goal box, in which they were confined for 30s. The affective valence of the object-cues for subsequent conditioning sessions was determined based on exploration latencies during habituation 3 & 4. The two most-explored object-cues were assigned as the aversive cue-pair, the two least-explored as the appetitive cue-pair, and the remaining two as the neutral cue-pair.

##### Object-cue conditioning

The animals were trained to associate the 3 object-pairs with either appetitive (sucrose), aversive (mild foot shock), or neutral (no event) outcomes over 9 daily conditioning sessions. Each rat completed 3 trials per day, 1 for each affective valence. During a trial, the time spent exploring each object-cue was recorded, and the rat could enter the neutral box after 1 min elapsed. After 10s, rats could enter the goal box where they were confined to for 30s during which the outcome assigned to the explored object-pair was administered (appetitive: 2 x 0.8 ml of 20% sucrose every 15s; aversive: 2 x 0.75s, 0.26-0.29 mA shock every 15s; neutral: no outcome). After 30s elapsed, rats were removed from the apparatus and returned to their home-cage in preparation for the next conditioning trial. If rats did not readily enter the next box within 30s, they were gently ushered in by the experimenter, and the latency to enter (LTE) the object, neutral, and goal boxes was also recorded as a potential measure of preference/aversion of expected outcome. The shock level was calibrated for each rat during the first aversive conditioning session and fixed at a level which elicited a mild startle response and defensive treading behaviour, but not freezing. The order of presentation of the object-cues, and the order in which rats completed each trial were changed daily.

##### Conditioned cue acquisition test

Acquisition tests performed under extinction conditions were conducted after conditioning days 4 and 8 in order to assess learning of the object cue-outcome associations. The experimental procedure was identical to that of cue conditioning training, except that rats were not confined to the goal box upon entry and were permitted 2 min to freely move between the neutral box and goal box during this time. Successful acquisition was indicated by the rats spending more time in the goal box for appetitive trials than aversive and neutral trials, and rats spending more time in the neutral box for aversive trials than appetitive and neutral trials. Following the second acquisition test, rats were given a ‘refresher’ conditioning day prior to proceeding to the AA conflict test.

##### Approach-avoidance conflict test

On the day of AA conflict testing, rats were bilaterally tethered to the laser and placed into the start box. The laser was then turned on for animals completing the task under inactivation conditions, before the start box guillotine door was removed exposing the animals to the object box, to which they were confined upon entry. One appetitive object-cue and one aversive object-cue were presented in recombination as the ‘conflict object-pair’, which animals could freely explore for 1 min. Identical to conditioning sessions, rats were then given 30s to freely enter the neutral box, after which they were gently ushered in by the experimenter, and were confined here for 10s before being permitted to enter the outcome box. At this point, rats were given 3 min to freely move between the neutral box and outcome box, after which the laser was turned off and the animal was removed from the apparatus and returned to its home cage in preparation for the next trial. All rats completed two conflict test sessions, one with the laser on and one with the laser off for the entire duration of the test session, each with a different set of conflict object-cues. The number of entries into and ‘retreats’ from the outcome box were recorded; an entry was defined when the animal’s hind limbs stepped on to the grid floor of the outcome box whereas ‘retreats’ were defined as partial entry into the outcome box followed by an immediate exit or backward treading into the neutral box. The order of the laser treatment, the assigned combination of the conflict object-cues and whether the aversive or appetitive object was presented first in the runway were all counterbalanced.

##### Control tests

To ensure that PRC-inhibited could still discriminate stimuli that they were trained on during the conditioning session, a test with the neutral object-pair was conducted on the same day as the 2 AA conflict tests, during which half of the rats [PRC *n* = 9 (*n* = 5 ArchT); vCA3 *n* = 8 (*n* = 5 ArchT)] completed the test with the laser on, and half with the laser off.

To investigate whether reducing the level of object-associated motivational conflict might change PRC-mediated behaviour, both PRC- and vCA3-rats were given a ‘refresher’ conditioning session prior to administration of a ‘low conflict’ recombination test, during which either an appetitive or aversive object-cue was paired with a neutral object-cue. PRC- and vCA3-rats were each given two trials per test, an appetitive-neutral object-pairing, and an aversive-neutral pairing. Each rat was tested twice, once with the laser on throughout the entire test session, and once with the laser off. The order of laser treatment, the assigned combination of the object-cues, and whether the appetitive/aversive or neutral object was presented first was counterbalanced within and between each test session.

Finally, to control for the novelty of the recombined object-pair presented during both the ‘high’ and ‘low’ conflict tests, along with an additional measure to confirm that PRC-rats could still discriminate stimuli under optogenetic inhibition, all rats completed a ‘no conflict’ test, during which a novel recombination of objects within the same valence were presented. Rats were first trained to associate a novel set of 8 objects with either appetitive (4 objects; 2 object-pairs) or aversive outcome across 4 daily conditioning sessions, followed by an acquisition test and a ‘refresher’ conditioning session prior to testing. On the day of testing, rats [PRC *n* = 12 (*n* = 8 ArchT); vCA3 *n* = 8 (*n* = 5 ArchT)] were given two trials per test, an appetitive-appetitive object-pairing, and an aversive-aversive pairing. Each rat was tested twice, once with the laser on throughout the entire test session, and once with the laser off.

#### Object-based Approach-Avoidance Shuttle box Task

##### Habituation

All rats were given 3 days of habituation the runway apparatus. On day 1, rats could freely explore both sides of the apparatus for 5 min, with the central guillotine door raised. On day 2, the rats were confined to one side of the apparatus with the guillotine door and following 2.5 min of exploration, the door was raised and animals were permitted to shuttle to the opposite side of the apparatus. Upon entry, the door was lowered and the animals were confined for a further 2.5 min. On day 3, the rats were confined to one side of the apparatus which now contained two sets of removable panels – one opaque and one transparent, preventing access to the sucrose dishes located at the end of the two compartments. After 10s the opaque panel was removed, providing visual but not tactile access to the sucrose dispenser. After a further 30s, the transparent panel was removed and the central guillotine door was simultaneously raised. The animals could then freely explore the empty sucrose dish, and were permitted to shuttle to the opposite side, which contained another set of opaque and transparent panels. Once animals shuttled to the opposite side, the door was lowered, and the habituation procedure was repeated in the other compartment.

##### Object-cue conditioning

The animals were trained to associate the 2 object-pairs with either appetitive (sucrose) or aversive (mild foot shock) outcomes over 9 daily conditioning sessions. Each rat completed 4 trials per day, 2 for each affective valence. During a trial, rats were placed between the central guillotine door and the opaque panel, confining them to one side of the apparatus. Following 10s the opaque panel was removed and the animals were permitted to visually sample the object pair for 1 min, which remained behind the transparent panel. After 1 min elapsed, the transparent barrier was removed and the central door was simultaneously raised and outcome was immediately provided (appetitive: 1 x 1.5 ml of 20% sucrose; aversive: 1s 0.24-0.27 mA shock, pulsed every 2s). Animals were given 1 min to shuttle to the opposite side of the apparatus before they were gently ushered in by the experimenter. Upon shuttling, the central door was lowered and the next trial began. The time spent visually sampling the objects and escape latency data were collected. The order of presentation of the object-cues, and the order in which rats completed each trial were changed daily.

##### Conditioned cue acquisition test

Acquisition tests performed under extinction conditions were administered after conditioning days 4 and 8 in order to assess learning of the object cue-outcome associations. The experimental procedure was identical to that of cue conditioning training, except that the central guillotine door was not lowered upon the first shuttle response, and rats were permitted to re-enter the cued side of the apparatus and were given 3 min to freely move between the cued and uncued sides. Similar to conditioning sessions, the uncued side contained an opaque panel. Successful acquisition was indicated by the rats exhibiting significantly shorter escape latencies for aversive trials than appetitive trials, and spending less time in the cued side during aversive trials than appetitive trials. Following the second acquisition test, rats were given a ‘refresher’ conditioning day prior to proceeding to the AA conflict test.

##### Approach-avoidance conflict test

On the day of AA conflict testing, rats were bilaterally tethered to the laser and rats were placed between the central guillotine door and the opaque panel, confining them to one side of the apparatus. The laser was then turned on for animals completing the task under inactivation conditions before the opaque panel was removed exposing the animals to the object-pairs behind the transparent panel. One appetitive object-cue and one aversive object-cue were presented in recombination as the ‘conflict object-pair’, which animals could visually sample for 1 min. Once 1 min had elapsed, the transparent panel was removed and the central door was simultaneously raised. At this point, rats were given 3 min to freely move between the ‘conflict-cued’ side of the apparatus and the uncued side, after which the animal was gently ushered into and confined to the uncued side by the experimenter. The laser was then turned off for animals completing the subsequent conflict test without inactivation. All rats completed two conflict test sessions under extinction conditions, one with the laser on and one with the laser off for the entire duration of the test session, each with a different set of conflict object-cues. The escape latency and time spent in the ‘conflict-cued’ side of the apparatus were recorded, along with the number of re-entries into the cued side; a re-entry was defined when the animal’s hind limbs crossed the threshold of the central door of the cued side. The order of the laser treatment and the assigned combination of the conflict object-cues were all counterbalanced.

##### Control tests

Following the conflict test, animals were given a ‘refresher conditioning’ session, prior to completing 2 sets of conditioned cue acquisition tests, one with the laser on and one with the laser off. The procedure of the tests was identical to the conditioned cue acquisition test described above, which were conducted under extinction conditions. On the day of testing, the laser was turned on for animals completing the task under inactivation conditions before completing 1 appetitive and 1 aversive acquisition test. On the next day all rats were given another ‘refresher conditioning’ session before completing another set of acquisition tests the following day, with the laser turned off for the animals completing the task without inactivation conditions.

#### Radial Arm Maze Task

##### Habituation

PRC-rats underwent four, 5 min habituation sessions as previously described (Nguyen et al., 2019; Schumacher et al., 2018). Briefly, on day 1, following 1 min confinement to the central hub, all guillotine doors were opened allowing 5 min of free exploration of the three arms without cues present. On day 2, a different set of cue inserts were placed into each arm and rats could freely explore the cues for 5 min. The affective valence of the cues for subsequent conditioning sessions was determined based on exploration time during this session. The most-explored cue was assigned as the aversive cue, the least-explored as the appetitive cue, and the remaining one as the neutral cue. On day 3, two guillotine doors were lifted and rats could explore the newly assigned neutral cue in one arm, and a superimposition of the appetitive and aversive cues in the other arm for 5 min; this would mirror the conditions for the final conflict test. The final habituation session was done without cues and habituated rats to confinement within the arms. Following 1 min in the hub, the rats would sequentially enter each arm, and were confined to them for 1 min before being allowed to return to the central hub.

##### Cue conditioning

PRC-rats were trained to associate 3 sets of visuotactile cues with appetitive (sucrose), aversive (shock), or neutral (no event) outcomes over 9 daily conditioning sessions. The rat was placed in the central hub for 30s after which one guillotine door was raised permitting entry to an arm, followed by confinement in the arm for 2 min. During this time the outcome assigned to the cue was administered (appetitive: 4 x 0.4 ml of 20% sucrose administered every 30s; aversive: 4 x 0.75s, 0.26-0.29 mA shock delivered at a random point every 30s; neutral: no outcome). After 2 min elapsed, the door was raised and rats were permitted to return to the central hub, and this process was repeated for the remaining two arms. The assignment of each cue to a given arm and the order in which each cue and arm were presented was changed daily. While the relative shape of the maze was held constant between sessions (Y-maze configuration), the maze was rotated either clockwise or counter-clockwise by 60° for each training session to prevent the use of spatial cues.

##### Conditioned cue acquisition test

Acquisition tests performed under extinction conditions were conducted after the fourth and eighth training sessions. The test procedure was identical to day 2 of habituation in that rats were given 5 min to explore all 3 arms simultaneously, but with cue inserts under extinction conditions. Successful acquisition of conditioned behaviour was indicated by the rats spending more time in the appetitive arm than aversive and neutral arms, and rats spending less time in the aversive arm than the appetitive and neutral arms. Following the second acquisition test, rats were given a ‘refresher’ conditioning session prior to proceeding to the AA conflict test.

##### Approach-avoidance conflict test

The procedure for this test was identical to day 3 of habituation, with rats being permitted 3 min under extinction conditions to explore two arms presented simultaneously, one containing the neutral cue and one containing a superimposition of the appetitive and aversive cues. In addition to the time spent in each arm, the number of entries and ‘retreats’ into both arms were also recorded. All rats completed two test sessions on the same day, one with the laser on for the entire 3 min test period, and one with the laser off. The order of laser treatment and the assignment of the ‘conflict’ and neutral cues to a given arm were counterbalanced within and between test sessions.

#### Laser delivery

532nm (green) laser light was continuously applied for the length of time specified in each experiment. Laser illumination was delivered to an implanted optic fiber attached with plastic sleeves (diameter 2.5 mm; Doric Lenses, QC, Canada) via two steel-braided fiber optic cables (200µm core; 0.22NA; Doric Lenses), which received light through a bifurcating rotary joint (Doric Lenses) secured with an FC connector. The rotary joint was connected to a class 3-B diode-pumped solid-state (DPSS) laser (165mW output; Laserglow Technologies, ON, Canada) via an FC connector. The light output of the optic fiber was adjusted to approximately 15mW, and based on previous measurements (Deisseroth, 2012) incorporating geometric loss of light, this would produce an irradiance of 15.38mW/mm^2^ (0.22NA; fiber core radius = 200 µm; implanted 0.5mm above target site), which exceeds the minimum amount needed to produce opsin activation (Gradinaru et al., 2009; Stefanik et al., 2013; Tye et al., 2011).

#### cFos activation and histology

Prior to sacrifice, rats either completed a NOR task (PRC *n* = 18) or an elevated plus maze test (EPM; vCA3 *n* = 14) in order to endogenously increase c-Fos labelling in the respective brain areas. Previous work has demonstrated that c-Fos is consistently increased in the rat PRC following exposure to novel visual stimuli (Albasser et al., 2010; Zhu et al., 1995). Similarly, rats exposed to anxiety-provoking environments (e.g., EPM) consistently demonstrate increased c-Fos labelling in the vHPC (Duncan et al., 1996; Hale et al., 2008; Linden et al., 2004).

The NOR task was conducted in the open field apparatus (50 cm (L) X 50 cm (W) X 50 cm (H)]. Rats were first given 2 daily 5 min habituation sessions of the apparatus, without any object-stimuli. On the day of testing, the bedding in the apparatus was changed, and rats were placed in the open-field facing away from 2 identical objects (A1 & A2), which they were permitted to ‘sample’ for 5 min. Rats were subsequently returned to their home-cage for a 15 min delay interval. During the test phase, rats were returned to the open field, which now contained a third identical copy of the sample object (A3) and a novel object (B1), which they were permitted to explore for 3 min. Between the sample and test phases, the apparatus was not cleaned, nor the bedding agitated, and no other rats were placed in the apparatus during this time. Half of the rats received laser treatment [*n* = 9 (ArchT = 5)] during the sample and test phases, and half completed the task with the laser off.

The elevated plus maze apparatus consisted of a central area (10 cm (L) X 10 cm (W)) with four maze arms (43.2 (L) X 10 (W) X 43.2 (H)) forming a plus shape. Two maze arms, directly across from one another, were enclosed by high walls (24.8 cm (H)), while the other two arms were unenclosed or ‘open’. Rats were placed into the central area facing the ‘open’ arm, and were given 10 min to freely explore the apparatus. Half of the rats (*n* = 7 (ArchT = 4)) completed the task with the laser on, while the other half with the laser off.

Ninety minutes after completing the test phase in the NOR task or the EPM test, rats were administered a terminal dose of sodium pentobarbital (Bimeda-MTC, Cambridge, ON) and were intracardially perfused with phosphate-buffered saline (PBS), followed by 4% paraformaldehyde (PFA). Brains were removed and stored in 4% PFA for 24 h before sectioning. 50 µm coronal sections were obtained with a vibratome (VT1000s; Leica Microsystems, Germany), mounted onto glass slides with a Flouroshield mounting medium containing DAPI (Ab104139; Abcam). Optic fiber placement and viral injection site targeting were confirmed at 10x and 20x magnification under an Eclipse Ni-U epifluorescence microscope (Nikon Instruments, Japan) using a FITC filter.

For cFos immunoreactivity, sections were washed (5 times for 5 min with PBS on a shaker), and incubated in 1% H_2_O_2_ in PBS for 30 min. Sections were then incubated in 0.5% TNB blocking buffer for 1 h, followed by rabbit anti-cFos (in 0.5% TNB; 1:5000; Synaptic Systems), and left overnight at 4°C on a shaker. The next day, sections were first incubated for 1 h in donkey anti-rabbit (conjugated with horseradish peroxidase in 0.5% TNB; 1:500; Jackson ImmunoResearch), and then in diluted Rhodamine tyramide signal amplification (TSA) solution (in 0.01% H_2_O_2_ in 0.1M borate buffer; 1:500; Jackson ImmunoResearch) and wrapped in foil to prevent photobleaching for 30 min. Sections were mounted onto glass slides and treated with a mounting medium (Flouroshield, Ab104139; Abcam). Labelling of c-Fos proteins in the PRC and vCA3 were confirmed at 10x magnification under the Eclipse Ni-U epifluorescence microscope (Nikon Instruments, Japan) using FITC and TexasRed filters, with cell counting performed by an automated counting software (Fiji; Schindelin et al., 2012).

## Data analysis

All behavioural testing was recorded and tracked using Noldus EthoVision XT (Noldus, Netherlands). Data were analyzed using SPSS statistical package version 26.0 (IBM, ON, Canada). For the object-based AA task, conditioned cue acquisition test data for the time spent in the goal box, the LTE, number of entries and retreats, and object exploration latencies were analyzed by separate 3 x 2 repeated-measures ANOVA with a within-subjects factor of valence (appetitive, aversive, neutral), and a between-subjects factor of group. Analyzed data for the high-conflict test were identical to the acquisition test, and were subject to a 2 x 2 repeated measures ANOVA with within-subjects factors of laser treatment, and a between-subjects factor of group. Each rat only completed one session of the neutral test, with between-subjects factors of laser treatment and group, and data were analyzed by univariate ANOVA. Data for the low-conflict and no-conflict control tests were each first analyzed by separate 2 x 2 x 2 repeated-measures ANOVA with within-subjects factors of valence and laser treatment, and a between-subjects factor of group. Activity analysis by the sucrose dispenser and generation of heat map figures were completed using Noldus EthoVision XT, in which an 18cm (L) X 11.5cm (W) region of interest was demarcated within the goal box (around the dispenser; 1/3 of the total goal box), and could reliably contain an entire rat.

For the RAM task, conditioned cue acquisition data for the time spent in each of the cued arms, as well as the number of entries and retreats were analyzed by a 3 x 2 repeated-measures ANOVA with a within-subjects factor of valence (appetitive, aversive, neutral), and a between-subjects factor of group. Analyzed data for the conflict test were identical to the acquisition test, and were subject to a 2 x 2 x 2 repeated measures ANOVA with within-subjects factors of laser treatment and trial type (conflict vs neutral), and a between-subjects factor of group. For the EPM test data, 2 x 2 repeated-measure ANOVAs were conducted to compare the time spent in the open and closed arms, as well as the number of entries made into both, with a within-subjects factor of arm type (open vs closed), and between-subjects factors of laser treatment and group.

For the NOR task, a 2 x 2 repeated measures ANOVA was conducted on the ‘discrimination ratio’ (d_2_ 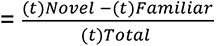), with a within-subjects factor of laser treatment and between-subjects factor of group. Behavioural data from the NOR task revealed that consistent with previous findings (Dix and Aggleton, 1999; Winters et al., 2004; Winters and Bussey, 2005), both novel object exploration and novel object discrimination performance were significantly higher during the first minute of the test session (time point: F(2, 28) = 20.79, p < 0.001; object x time point: F(2, 28) = 16.41, p < 0.001), and thus, this time-point was selected for subsequent analysis.

Significant main effects and interactions were further investigated with simple contrasts and post hoc tests with a Bonferroni correction where appropriate. Violations to sphericity were addressed with a Huynh-Feldt correction.

## Acknowledgements

This work was funded by the Canadian Institutes of Health Research (#156070 to A.C.H.L. and R.I.). The authors gratefully acknowledge Dr Maithe Arruda-Carvalho and the members of her DevNeuro lab in allowing us to use their Ni-U epifluorescence microscope.

## Competing Interests

The authors declare no competing interests.

---

**Supplementary Figure 1.**
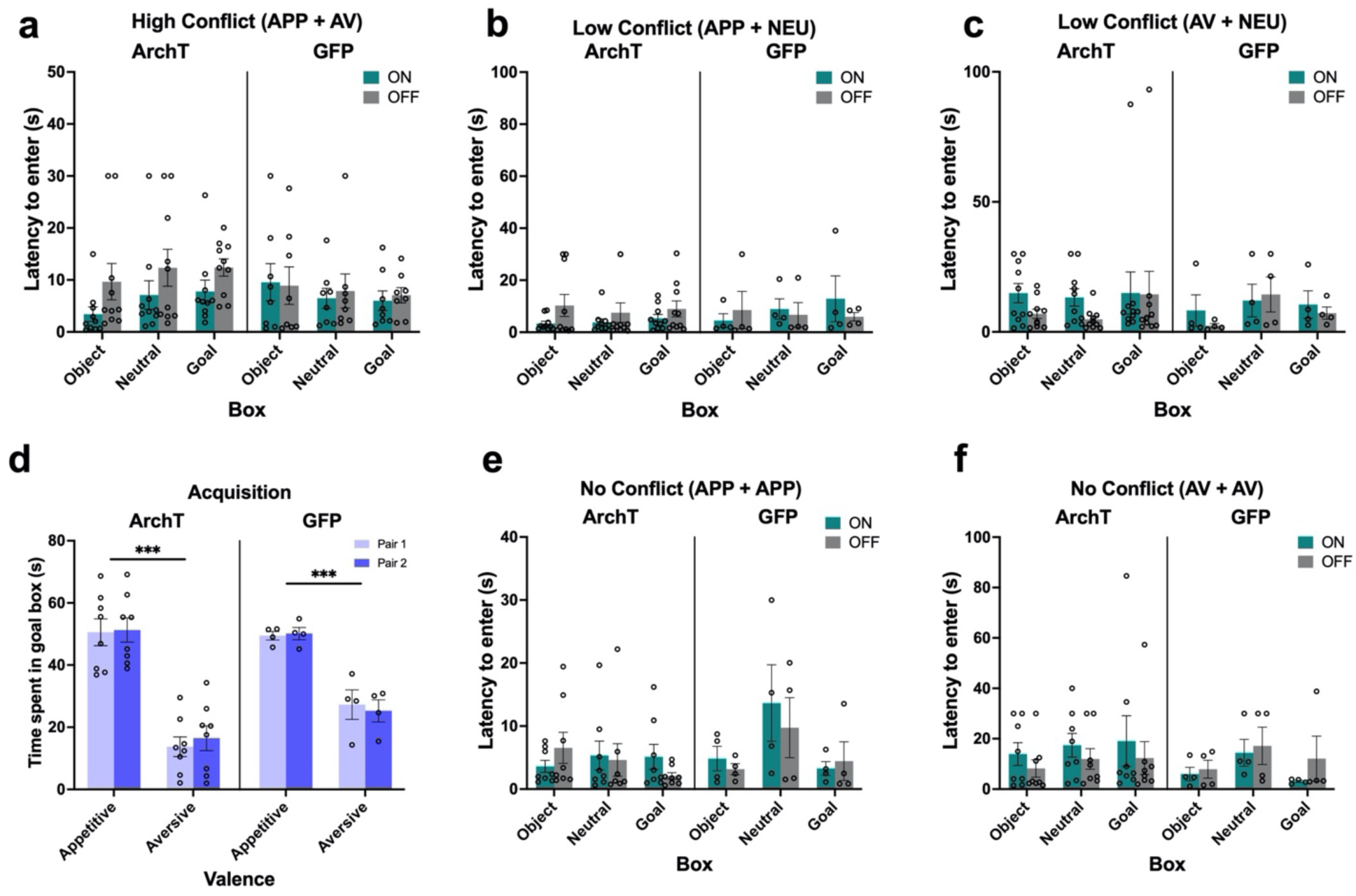
Additional PRC rat data for the AA conflict runway task. **(a-c)** PRC inhibition did not impact rats’ latency to enter the goal box in the high conflict test or each of the two low conflict tests. **(b)** Both ArchT and GFP PRC rats successfully learned a new set of appetitive and aversive object pairs for the no conflict recombination tests. **(e-f)** PRC inhibition had no effect on latency to enter in the appetitive or aversive no conflict recombination tests. *** p < 0.001

## Notes

### Competing Interest Statement

The authors have declared no competing interest.

